# Bioelectrical Synchronisation of Spruce Trees during a Solar Eclipse

**DOI:** 10.1101/2023.10.24.563666

**Authors:** Alessandro Chiolerio, Monica Gagliano, Silvio Pilia, Paolo Pilia, Giuseppe Vitiello, Mohammad Mahdi Dehshibi, Andrew Adamatzky

**Affiliations:** Bioinspired Soft Robotics, Istituto Italiano di Tecnologia, Via Morego 30, 16163 Genova, Italy; University of the West of England, Coldharbour Lane, Bristol, United Kingdom; The Cyberforest Experiment, Paneveggio (TN), Italy; The Biological Intelligence Lab, Southern Cross University, Lismore (NSW), Australia; Openazienda S.r.l., Macomer (NU), Italy; Dipartimento di Fisica “E.R. Caianiello”, Università di Salerno, 84084 Fisciano (SA), Italy; Departamento de Informática, Universidad Carlos III de Madrid, Leganés, Spain

## Abstract

Regular light-dark cycles greatly affect organisms, and events like eclipses induce distinctive physiological and behavioural shifts. While well-documented in animals, plant behaviour during eclipses remains largely unexplored. Here we monitored multiple spruce trees to assess their individual and collective bioelectrical responses to a solar eclipse. Trees anticipated the eclipse, synchronising their individual bioelectrical behaviour hours in advance. Older trees displayed greater anticipatory behaviour with early time-asymmetry and entropy increases, characteristic of memory dynamics in open dissipative systems such as trees. These results reveal a complex relationship between individual and collective tree behaviour, shaped by individual age and physiology as well as collective history and memory. This highlights the adaptive significance of synchrony and collective memory in plants, offering new insights on coordinated behaviours in nature.

**One Sentence Summary:** Trees anticipate and respond collectively to a solar eclipse.

## 1 Main Text

Sunlight and its periodicity drive global weather patterns, seasons and climate, and make life possible on our planet. Daily and seasonal cycles of natural light organise biological systems by synchronising their internal clock with the geophysical cycles of the Earth (*1*); disturbances to these rhythms initiate distinctive changes in the physiological and behavioural function of living organisms, exerting a strong constraint in terms of their adaptability and resilience to environmental changes (*2*). At a time marked by growing human-induced changes to natural cycles, unusual astronomical events such as eclipses effectively function as natural experiments, providing valuable insights into how living organisms respond to sudden and unanticipated changes in their environment (*3, 4*). Even with their infrequent and momentary occurrence, eclipses and their impact on organismal behaviour have been observed for millennia and are now well documented’ (*5*). In humans, for example, solar eclipses have played a transformative role inspiring awe and arousing social cohesion and prosocial tendencies, even capable of ending wars (*6*). Heightened tendencies of individuals to form collectives by huddling, gathering and synchronising group movements during solar eclipses are also seen across several animal groups, terrestrial and aquatic alike (*7*). If solar eclipses play such a vital role in shaping individuals and their groups to ensure collective survival across species, it is remarkable that very little is known about how plants respond to these astronomical events and such knowledge is limited to responses at the individual level (*8–14*) overlooking that group behaviour has also been observed in plants (*15, 16*). Here, we use a remote measurement system we have developed to simultaneously monitor multiple trees in a forest (*17*) to directly test whether individual trees respond to a solar eclipse together as a larger living collective, the forest.

To investigate this possibility, we examined the electrome (*18*) of spruce trees (Picea abies) to describe the complexity of their bioelectrical activity during a partial solar eclipse that occurred in a forest located in the Dolomites mountain region, north-eastern Italy (Fig. 1 A and B). Electrome dynamics emerge from a blend of electrical signals plants use to coordinate an assortment of physiological functions necessary to readily respond to environmental fluctuations (*19, 20*). These dynamics are known to be non-random, exhibit long-range temporal correlations and persistence (*21*), and are most likely associated with information content and transduction in long-distance signalling (*19, 22, 23*). In a previous study conducted in this forest, we had established a methodological framework for measuring, sorting, and analysing the electrome dynamics of spruce trees at different developmental stages (i.e., healthy and young, healthy and old, dead logs) and determined that tree electromes correlated with solar (and lunar) cycles (*17*). Therefore, in this study, we hypothesise that the different phases of a behavioural response to a specific event, such as a solar eclipse, can be detected by monitoring how the electrome changes as the eclipse occurs.

**Figure 1:**
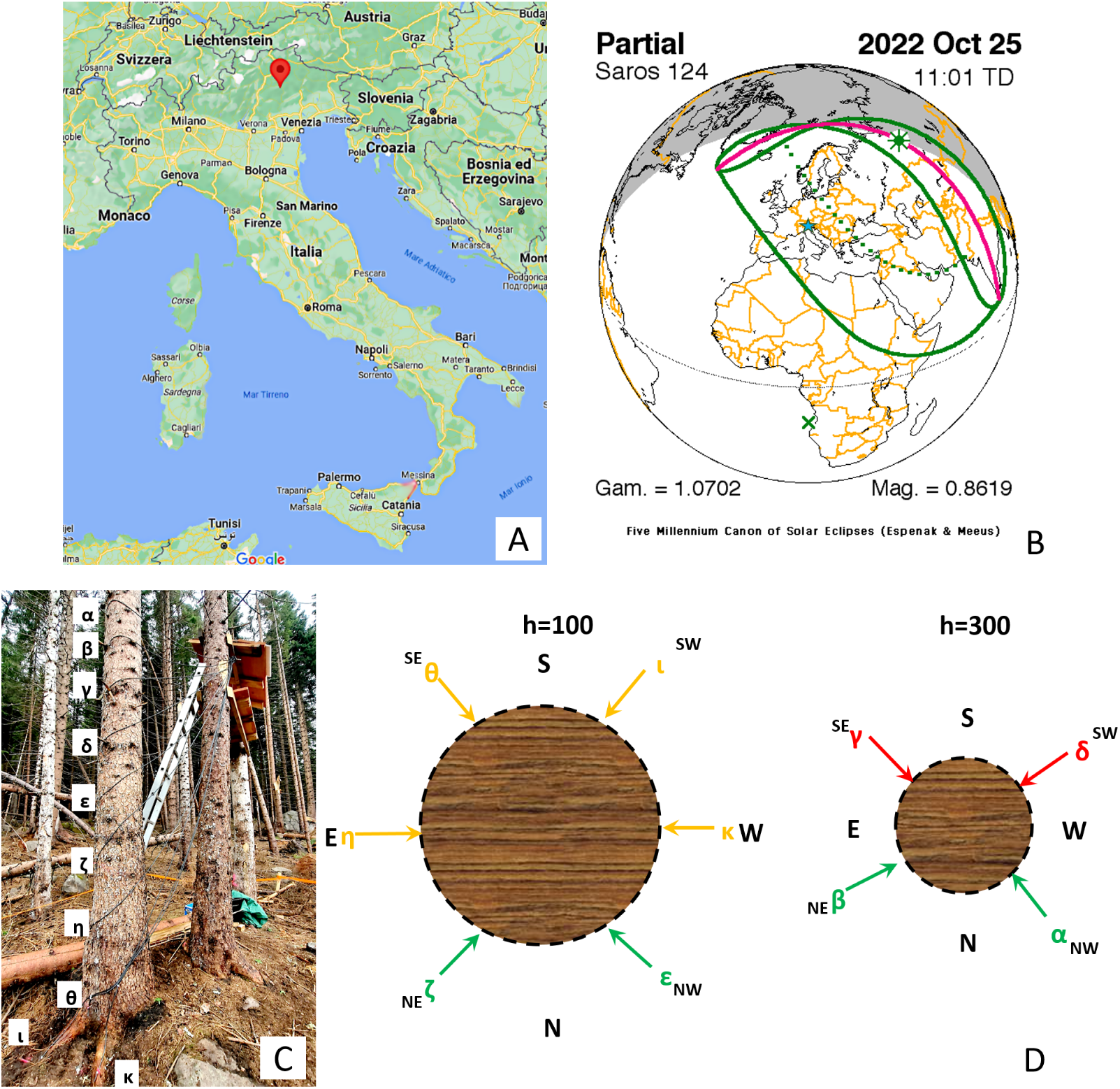
Experimental set-up to simultaneously monitor the electrome of multiple trees during a solar eclipse. (A) The location of the experimental site at the Costa Bocche forest near Paneveggio in Dolomites area, Italy. (B) Diagram of the Saros 124 event, the solar eclipse that occurred on 25*^th^* October 2022. Green continuous lines trace the Sun shadow at ground level, the dashed line corresponds to a coverage of 50% of the Sun disk, the pink line to the Sun path. (C) In-situ installation of 10 xilematic electrodes (labelled with the Greek letters) on a spruce tree. Stainless steel threaded rods of 6 mm diameter were spaced at 50 cm apart along the trunk and in contact with the core of the tree. Each electrode was then connected to the Cybertree devices via low impedance audio cables. (D) Horizontal view of the phloematic configuration with electrodes in contact with the superficial layer of the tree and radially arranged in two arrays. (i) One array was located at 1 m above ground with electrodes at a radial distance of 60° from each other; (ii) the other array was located at 3 m above ground with electrodes at a radial distance of 60° from each other. Cardinal and ordinal directions are indicated.

We monitored and recorded the bioelectrical activity of spruce trees using a network of CyberTree devices; these are low-cost, low-power consumption, low-temperature resistant devices custom-built to provide remote readings of tree electrical signals (as described in the Supplementary Materials, Table S1-S3, Fig. S1-S6). The CyberTree devices enabled us to simultaneously collect data from multiple individual trees (n=3 healthy trees and n=5 tree stumps) from four sites (D, E, F and G) within the same forest (Figure 1 C). Specifically, we compared the bioelectrical activity of two healthy 70-year-old trees, one positioned in full sun at site D, and one in full shade at site F; one healthy 20-year-old in full shade at site G, and five stumps of old pristine trees at site E, which had been devastated by the storm Vaia in 2018. For each tree, we collected the bioelectrical potentials from five differential electrode pairs, each electrode pair being connected to a differential amplifier prior to recording the data (see details in Supplementary Materials). The electrodes were labelled with the Greek letters *α, β, γ, δ, ɛ, η, ζ, θ, ι, κ*. The set-up can collect information from both the core of the tree (xilematic), and its outer layers (phloematic), as shown in Fig. 1 D. We analysed the bioelectrical responses of these trees during the partial solar eclipse using higher-order complexity measures. As in similar linear/nonlinear adaptive signal-processing systems, first-order statistics, such as mean, variance, and correlation, become inadequate when the data deviate from a Gaussian distribution and the adaptive system exhibits nonlinearity (*24*). Further, the concept of information theory suggests that the level of uncertainty determines the information value of data (*25*). Indeed, the data carry little information if an event is highly probable. Using metrics, such as the entropy, Simpson diversity, expressiveness, Lempel–Ziv complexity, and fractal dimension, higher-order measures help to reduce uncertainty (*17*). Therefore, we computed eleven information-theoretic complexity measures to characterise the spatiotemporal activity patterns in raw data within a 10-minute time window (detailed in Supplementary Materials).

### 1.1 Individual tree responses to the solar eclipse

Based on the starting and ending point of the solar eclipse, we detected a general trend directly associated with the geometric occultation of the Sun and corresponding to an overall reduction of the bioelectrical potential at all sites with living trees. At the shaded site G, for example, the young 20-year-old tree exhibited a reduction of its bioelectrical potential, evident particularly for the electrode pairs *ɛ − ζ* and *η − θ* (Fig. 2A). Remarkably, we detected a sharp spiking activity, particularly at the levels of the two upper pairs *α − β* and *γ − δ* and visible as several quasi-square-waves concentrating until the onset of the eclipse, and then disappearing in the phase that immediately follows the end of the eclipse (Fig. 2 B-D).

**Figure 2:**
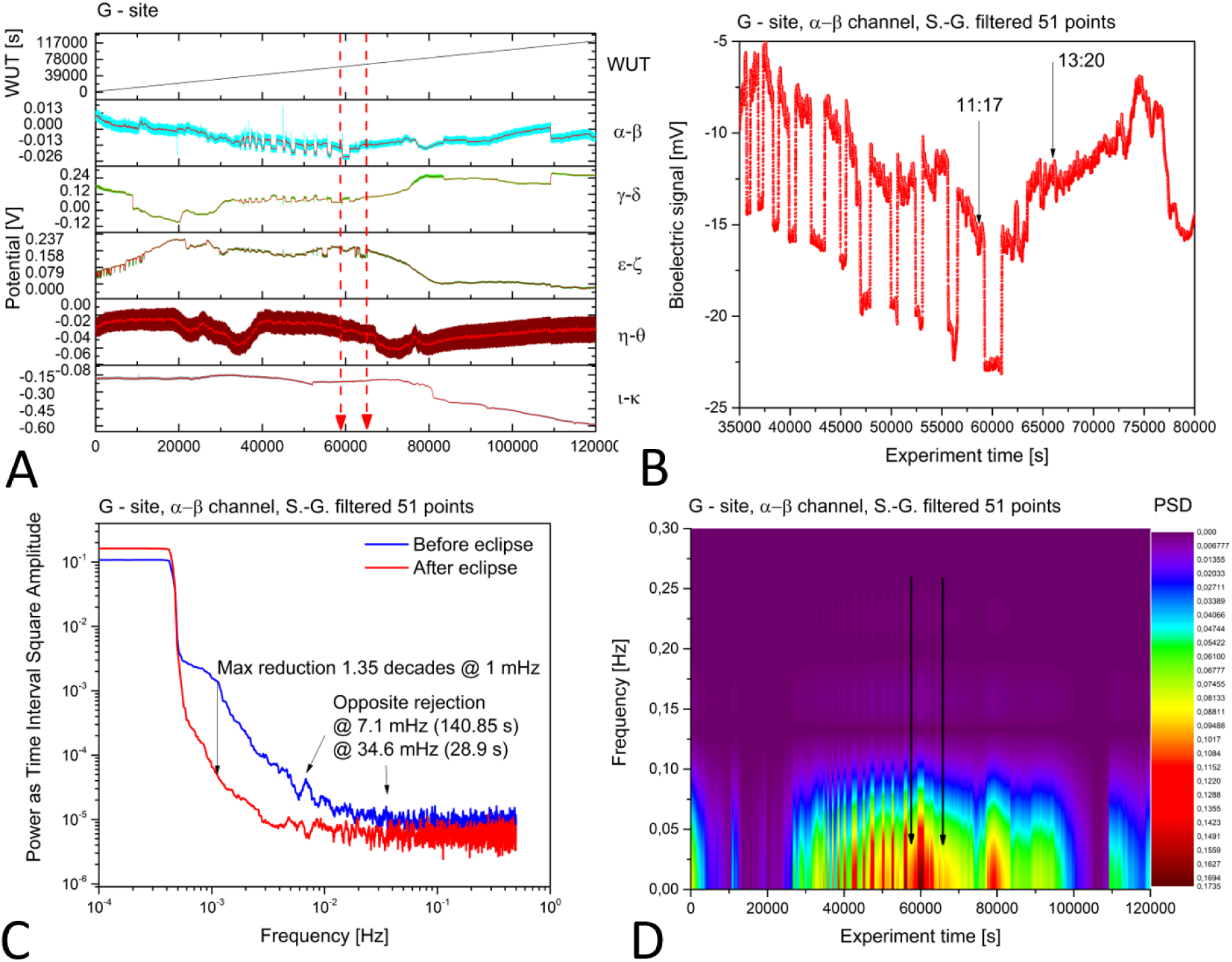
Bioelectric potentials recorded from the 20-year-old tree at site G during the solar eclipse. (A) Raw measurements collected from five differential channels. Data chunk spanned 120000 s and featured the five differential channels (curves) plus the “wake-up time” (WUT) of the CyberTree device as a reference to correlate with spikes that might occur during eventual restart. In this specific case the WUT corresponds to the experiment duration. The two red arrowed broken lines indicate the starting (11:17) and the ending (13:20) of the solar eclipse. (B) Enlarged view of the *α − β* differential channel with solar eclipse timing indicated. A first-order Savitzki-Golay (S.-G.) function with 51 points was applied to smooth the data and allow the underlying oscillations and correlations between the five differential channels to become more noticeable. (C) FFT analysis of 40.000 s before the solar eclipse and 40.000 s after, taken from the *α − β* differential channel. S.-G. smoothing applied. As indicated, the noise power dropped by 1.35 decades @ 1 mHz, and two peaks showing opposite rejection were found around 7 and 35 mHz, corresponding to periods of 141 and 29 s respectively (D) STFT analysis of the entire data shown in panel (A), limited to *α − β* differential channel. The black arrows indicate the starting and the ending of the solar eclipse. S.-G. smoothing applied.

A more in-depth analysis of the signals collected from channel *α−β* comparing all complexity measures allowed us to appreciate the noise and fluctuation of almost all functions before the eclipse, and a strong reduction afterwards (Fig. 3). For example, the expressiveness stopped its 5/10 minutes oscillating behaviour at the beginning of the eclipse, went into a smoother period that lasted hours after the end of the event, and then recovered to the original absolute values (Fig. 3 B). Similarly, the fractal index showed a strong oscillation until the beginning of the eclipse, then a transition to a smoother area (Fig. 3 C), indicating that the bioelectric signal from the tree lost the natural variability in spiking and oscillations which is observed as spontaneous electrical activity during a normal day.

**Figure 3:**
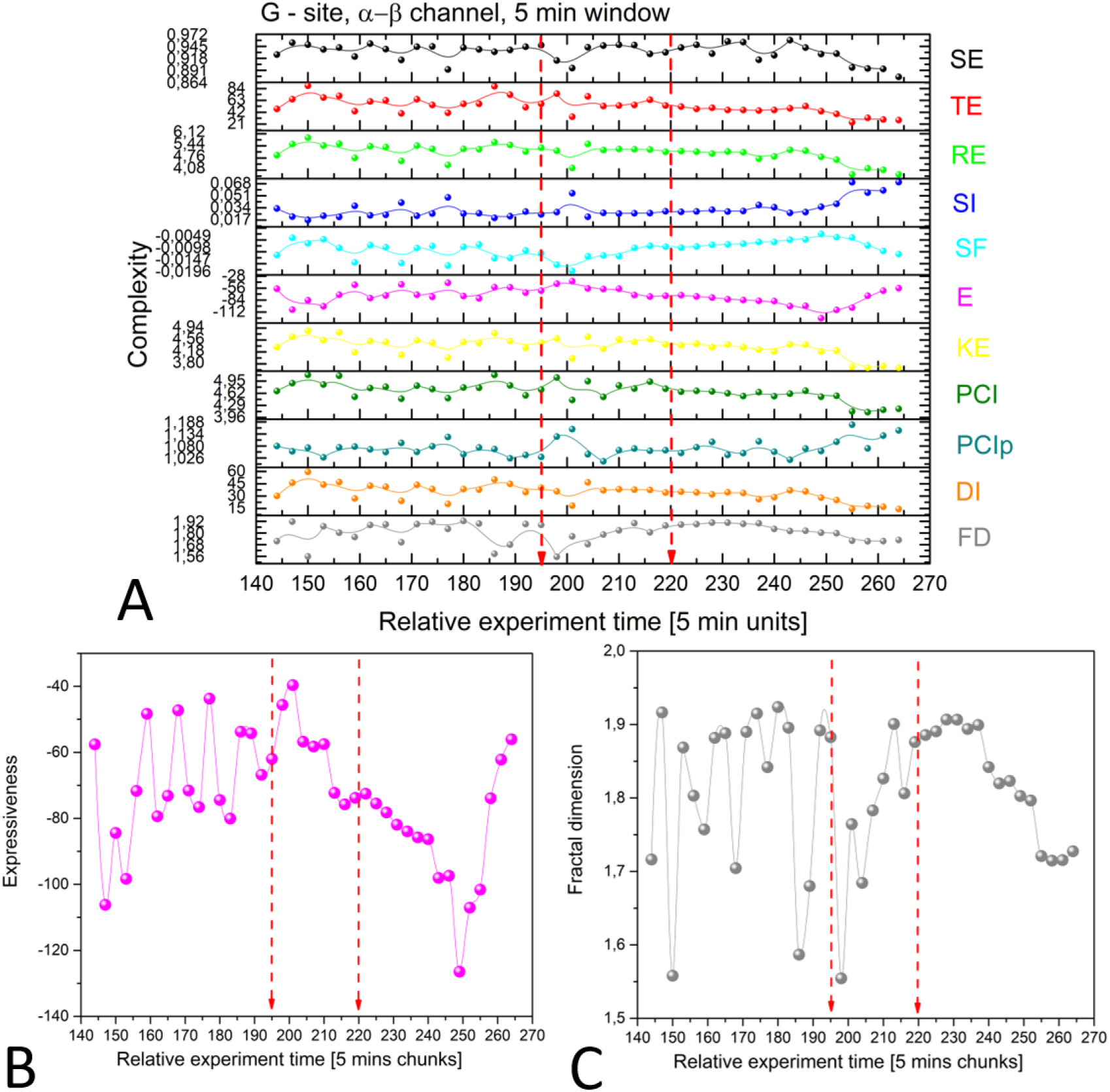
Complexity analysis of bioelectrical potentials recorded from the 20-year-old tree at site G during the solar eclipse (indicated by the two red arrowed broken lines). (A) Comparison of all the complexity measures using data chunks of 5 minutes: Shannon entropy, Tsallis entropy, Renyi entropy, Simpson index, Space-filling, Expressiveness, Kolmogorov entropy, Perturbation complexity index, permuted Perturbation complexity index, Diversity index, Fractal dimension. The two red arrowed broken lines indicate the starting (11:17) and the ending (13:20) of the solar eclipse. Detailed variation for (B) the Expressiveness and (C) the Fractal dimension during the solar eclipse.

At sites D and F, where we monitored the older trees, we also observed a reduction of the bioelectrical potential corresponding to the Sun occlusion (Fig. S7). On the F-site tree, this reduction was particularly sharp at the top electrode pair *γ − δ*, but also detected at the level of the three bottom electrode pairs *ɛ−ζ*, *η−θ*, and *ι−κ*, though taking place over a slightly broader timescale (Fig. S7 E). The D-site tree shows a bioelectric potential discontinuity, matching well the timing of the eclipse, on at least two electrode pairs, specifically the top one *α − β* and the bottom electrode pair *ι − κ* (Fig. S7 A). In these older trees, we detected a slow change in the initial bioelectrical potential level by approximately 300% occurring during the first 50000s (Fig. S7 E). For F-site tree, for example, this trend was evident in the bottom electrode pairs *ɛ − ζ*, *η − θ*, and *ι − κ*, where the pair facing the sun and oriented southwards (*ɛ − ζ*) detected increased bioelectrical potentials and higher noise level, while the other two pairs detected a reduction. This trend was not found in the top pairs *α − β* and *γ − δ*. From the end of the eclipse until 120000 seconds, all electrode pairs on the F-site tree detected a renormalization of the bioelectrical potential levels approaching their initial values. Lastly, we detected less pronounced variations in the bioelectrical responses of the five tree stumps at the E-site (Fig. S7 C). However, a deeper analysis based on FFT tools revealed a clear difference in the biopotential spectrum before and after the eclipse. We see that the oscillation frequency increases after the astronomical event. Being the xilematic electrodes connected more to the inner physiology of the plant, and to the movement of fluids inside the trunks triggered by capillarity and water pressure in the soil, we deduce that a higher frequency in the oscillation of liquids is generated. Oscillating masses reduce their fluctuation periods when the mass increases, therefore it is likely that the effective mass has increased.

### 1.2 Collective responses to the solar eclipse

To further investigate whether individual trees responded to the solar eclipse together as a collective, we conducted an analysis of stochastic processes in a multi-layered spatiotemporal arrangement (Fig. 4). Within each site, we observe the propagating bioelectrical waves, which can be attributed to various factors such as the movement of lymph carrying ionic currents (observed at D-site, where xylem electrodes are positioned along the trunk), circadian processes following the Sun’s orbit (seen at F-site, where phloem electrodes are arranged radially along the trunk), spontaneous spike-like processes (observed in the young tree located at the G-site), or even noise from recording logs (Fig. 4 A). Each stochastic process interacted with the others, akin to waves on the surface of water, exhibiting a certain level of temporal correlation (Fig. 4 B).

**Figure 4:**
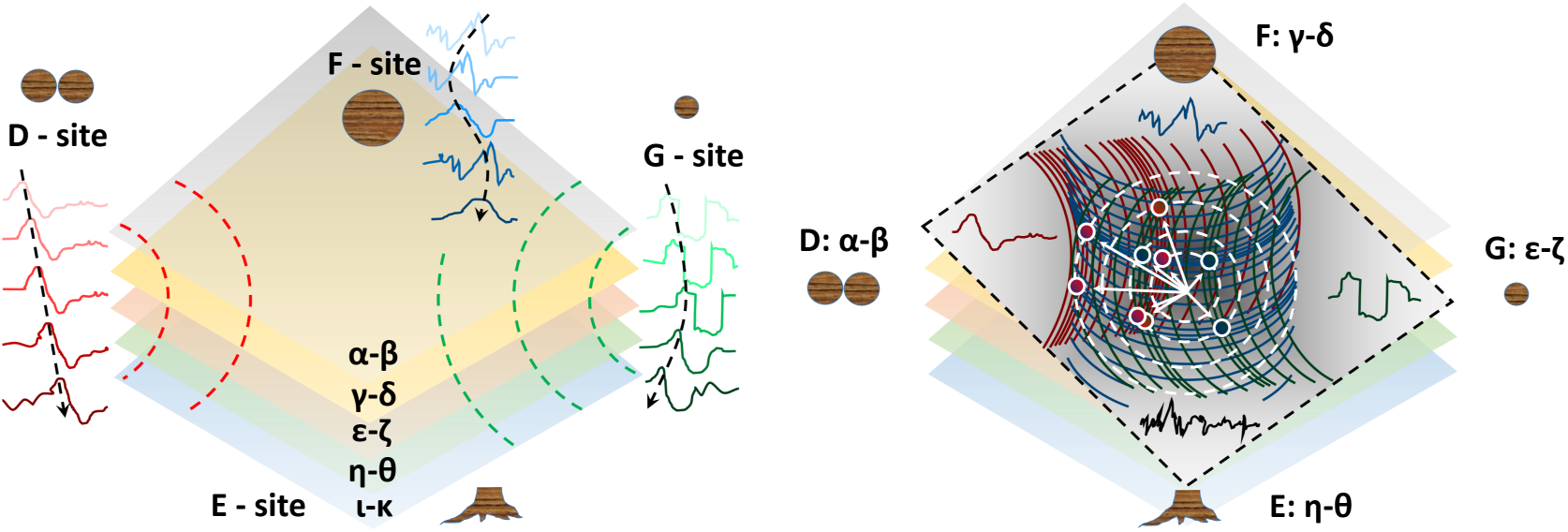
Bioelectrical potential recordings as stochastic processes in a multi-layered spatiotemporal arrangement. (A) Each corner of the square corresponds to a recording site (from left to bottom, in a clockwise direction: D-site, F-site, G-site, and E-site). Each set of recordings shows bioelectrical waves that propagate along the layers. (B) Interference of stochastic processes showing constructive/destructive profiles against a common barycenter (white arrows). Red waves propagate from the D-site, blue waves from the F-site, and green waves from the G-site. Constructive interference is pinpointed by white disks, whose inner colour gradient shows the fusion of the two originating processes.

By calculating the cross-correlation function, as described in Equation 1, we were able to determine the precise timing when two processes were most synchronised. In the equation, *g* represents the cross-correlation function, *d* and *f* denote two measured processes (bioelectric potentials), *r* symbolises the Cartesian volume in the space where the physical processes are collected, and *t* represents time.

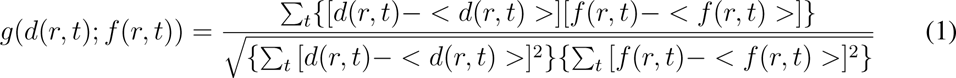

The cross-correlation functions calculated during a normal day exhibited a degree of symmetry with an absolute variation of around 0.1 units (Fig. 5 A). However, the degree of symmetry changed during the solar eclipse; absolute variations doubled in magnitude and had a smoother shape (Fig. 5 B). In other words, during a normal day, there was little difference between any two instants in the bioelectrical propagations occurring between two trees. During the eclipse, however, any two generic instants were indeed very different from each other. This interference was particularly noticeable between the trees D and F, which is unsurprising given that, because of their age, older trees serve as hubs that boast a more complex network of connections within a forest (*26*).

**Figure 5:**
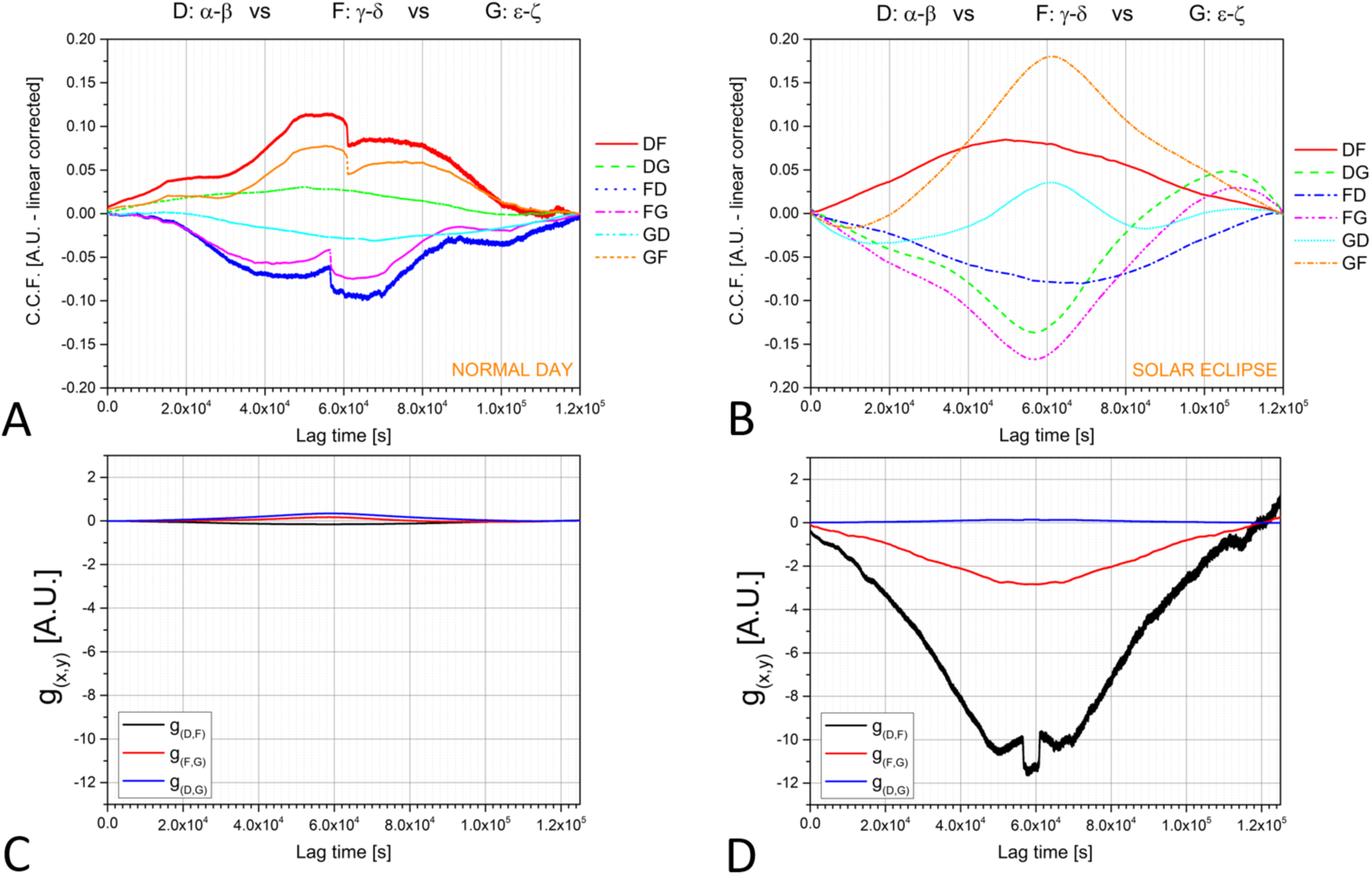
Cross-correlation functions calculated by comparing couple-wise processes occurring at the D-, F- and G-site (A) Cross-correlation functions calculated during a normal day. The choice of signals was totally random, comparing differential channels without any prior logic. Midday occurs at 6 *×* 10^4^ seconds. (B) Cross-correlation functions calculated during the solar eclipse. (C) Difference between direct and inverse cross-correlations during a normal day, and (D) during the solar eclipse.

The effects of the solar eclipse could also be monitored by calculating the difference between direct and inverse cross-correlations; for example, by considering the signal propagation from D- to F-site and from F- to D-site (black line in Fig. 5 C, D). As formalised in Equation 2, the differential function of implicit space-time variables *h* is a measure of the symmetry of two processes and should be zero for symmetrical ones:

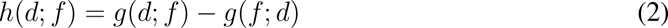

In this analysis, signal propagation was a temporal process. Therefore, we investigated whether a predictable event like the circadian cycle dictated by the daily rising and setting of the sun appears symmetric or ordinary when compared to a less frequent event such as the solar eclipse. By putting the two groups of complementary differences on the same scale, it is clear how the daily solar cycle produces very symmetric processes with deviation in time smaller than 1. When the forest experienced the solar eclipse, however, the effects were much more pronounced with a deviation from symmetry that could be greater than 10 with maximum values for the older trees (the D-F couple). Not only the deviation from temporal asymmetry was more pronounced in this couple of older trees, but the symmetry-breaking process (and the accompanying increase in entropy; (*27*) described by the vertical offset from zero, also commences earlier (by 1 hr) and concludes later than in couples with old and young trees (see the offset in the black curve, Fig. 5 D). Lastly, by comparing the time lags between simultaneous signals, we were able to conclude that the forest responded to the eclipse approximately 14 hours ahead from the actual occurrence of the event.

### 1.3 Theoretical modelling of collective bioelectrical activity

To understand what might be the origin of the time symmetry breaking observed in our experimental findings, we formulated in (*17*) and (*28*) a theoretical modelling based on quantum field theory (QFT), which we adopt also in the present discussion. The fundamental concept in the formulation is that trees are open, dissipative systems. The canonical formalism, however, only applies to closed systems, thus requiring the forest to be closed towards the environment. The closure is obtained by balancing the fluxes outgoing from the forest and incoming to the environment, and vice-versa. This is achieved, in the well known formalism of thermal QFT, called Thermo Field Dynamics (TFD), by doubling the tree’s degrees of freedom, say 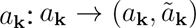, where 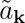 denote the environment degrees of freedom (see Supplementary Text for formal mathematical details). The state describing the ‘couples’ 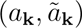 (i.e. the system and its environment) is a generalised coherent state (given by Eq. (S6)) (*29*). It is a thermal state, presenting entanglement between each tree and its environment, including therefore also other trees in the forest. The covariance of the state, providing a measure of the degree of entanglement, is non-zero (see details in Supplementary Materials). Based on previous observations of this system, the entanglement between the molecular activity of trees accounts for the observed non-vanishing value of the cross-correlation function in Eq. 1 (*17*). Also, the variations observed during the eclipse event (see Fig. 5 A, B) are due to the perturbations induced by the eclipse on the coherent condensate of the microscopic state at molecular and ionic level, largely dependent on the electric dipoles of the water molecules, resins, lymph molecular components, and other biomolecules of the tree (*17*). During the event, phase correlations among trees appear to be enhanced, which is mirrored by the condensate and, in turn, the bioelectric potentials. As trees are open systems and their evolution is ruled by dissipative dynamics (*28*), we expect non-symmetric reactions to perturbing agents, especially in older trees that have developed a larger number of correlations with the environment. As expressed by Eq. 2, such an asymmetry is expressed in the non-commutative character of the two processes *d* and *f* (see Fig. 5 C, D for other pairs of trees), hence indicating that the occurrence of the solar eclipse disrupts the symmetry of the communication channels shared by the trees. Remarkably, the theoretical modelling does indeed show that the anti-symmetric state is the lowest energy state, and therefore the one actually realised, as observed in our measurements (see formal details at the end of Section S0.5 of Supplementary Materials).

Being largely dependent on fluctuations of the electrical dipoles of a range of biomolecules, the tree microscopic dynamics is invariant under dipole rotational symmetry transformations. Several environmental influxes, including events such as an eclipse, disrupt such symmetry by selecting preferred directions in the molecular dipoles and associated ionic currents. Such conditions produce Spontaneous Symmetry Breakdown (SBS) (*30*), so that the ground state of the tree is no longer symmetric under the rotational symmetry of the dynamics. The molecular dipole fluctuations act as quantum variables and the SBS mechanism in QFT is known to generate long-range correlations among the system’s elementary constituents (the electric dipoles, in our case). The quanta associated with the long-range correlations in our case are called Dipole Wave Quanta (DWQ) and they coherently condense in the system ground state, which has then the structure of a coherent state. Here, the term coherence denotes that long-range correlations are “in phase”, hence coexisting in the ground state without reciprocal destructive interference.

Grounded on such a theoretical basis, our modelling then accounts for the observed behaviour of the cross-correlation function and several other features (see Fig. 5). For example, the electromagnetic (EM) vector potential **A**(**x***, t*), and therefore the bioelectric potentials from it derived, is directly affected by the coherent condensation of DWQ through the gauge transformation **A**(**x***, t*) *→* **A**(**x***, t*) + ***∇****λ*(**x***, t*), where the Coulomb gauge ***∇*** *·* **A**(**x***, t*) = 0 is assumed, and with it the constraint ***∇***^2^*λ*(**x***, t*) = 0, with *λ*(**x***, t*) related to the DWQ condensation density and therefore to the external agent triggering the SBS. The approach thus describes a direct relationship between the measured bioelectric potentials and environmental changes from atmospheric effects, including circadian cycles as well as light and gravitational effects before, during, and after the eclipse. The theoretical modelling also accounted for the observed fractal dimension features. In fact, it is known that there exists an isomorphism between deformed coherent states and fractal self-similarity (*31–33*). Changes in the fractal dimensions correspond to changes in the condensation densities of coherent states i.e. “deformation”, corresponding in turn to transitions between different dynamical regimes of the tree system. In the present case, for example, the smaller fractal dimension during the solar eclipse denotes a stronger coherent condensation density compared to the ones before and after the event (Fig. 3 C); in fact, the smaller fractal dimension signals a higher coherent regime. We conclude that the solar eclipse produces a SBS two orders of magnitude greater, and a consequent higher coherent condensation of DWQ, compared to normal daily cycles. This condition can be observed especially in older trees, where thicker xilematic tissues featuring richer molecular fluxes are present.

For the forest, the minimization of the free energy *F* is of course satisfied at each time t as: *dF* = *dU −* (1*/β*)*dS* = 0, i.e., *dU* = (1*/β*)*dS*, where *dU* and *dS* denote the variations of internal energy *U* and entropy *S*, and *β* = 1*/k_B_T* is the inverse temperature, with *k_B_* the Boltzmann constant. In terms of variations in the time 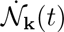 of the DWQ condensed number *N***_k_**(*t*), the free energy minimising condition leads to (*34*)

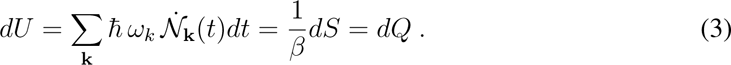

We see that as entropy decreases (and ordering increases), the rate of loss in time of coherent correlations decreases. As kinematical energy gets stored in the ordering of the system (*U* decreases), *dU → TdS*, we observed a power loss in the spectral analysis as the eclipse commenced and during the event (Fig. 2 C), and until the subsequent normal regime was reached after the eclipse (*dU ← TdS*). Once the event was no longer active, the forest reacted by releasing the energy stored in ordered patterns to the kinetic regime. A similar process aimed to restore the system to the state prior to the perturbing action was also observed in the fractal dimension behaviour (see Fig. 3 C and 5). By comparing direct and time-reversed cross-correlation functions of the bioelectric potentials, our modelling adequately corroborates our experimental findings, showing that the forest can be understood as a correlated complex system. This means that while the influence of the environment on the individual state and bioelectrical response of the trees is paramount, the biological correlations emerging as energy dynamically stored and released by the system as a whole, turn the trees into a single organism, the forest. By virtue of the collective sensing arising at the group level from synchronised individual responses, the forest reproduces the different phases of the individual’s responses to the changing conditions to respond to an event like the solar eclipse.

Synchrony plays a role in almost all aspects of collective behaviour and patterns of synchronous activity have been observed in almost every animal group studied (*35*). Notably, any synchronised behaviour fundamentally involves the synchronisation of movements (*36*). Hence, likely due to their sessile nature, behavioural synchronisation in plants has remained unexplored (and we are not concerned with phenological synchronisation on the well-known circadian or seasonal timescales, such as masting (*37*), flowering (*38*) or oscillatory motions of stems and leaves (*39*)). By monitoring the electrome of spruce trees in their natural environment during the eclipse, our field study provides evidence that trees synchronise their individual bioelectrical behaviour to generate a collective response to a shared source of external disturbance. Prior to this synchronised behaviour, the trees exhibited anticipation, a critical feature of synchronisation behaviour displayed by many animal species that makes synchrony distinguishable from repetitive reactions to repetitive stimulation (*36*). Specifically, the trees anticipated the solar eclipse by changing their bioelectrical behaviour several hours before the actual onset of the celestial event, thus preparing to cope with the perturbations the eclipse was expected to cause, such as substantial drops in sap flow rates within trees (*10*) and disruptions in their vital hydraulic connection between the soil and the atmosphere (*40*).

Crucially, organisms must be able to detect cues alerting them that circumstances have changed in value to ensure that the right adaptive response occurs at the right time through anticipatory behaviours (*41*). The cues that generated the anticipatory behaviour observed in these trees remain to be determined. We exclude sunlight and air temperature alterations induced by the solar eclipse as potential cues because the tree anticipated the event 14 hours prior to its occurrence, when both Sun and Moon were at their zenith on the other side of the Earth (i.e., on the Solomon Islands and off the coast of Vietnam, respectively). However, potential cues might have temporal patterns dependent on Sun-Moon-Earth orbital dynamics. We propose that the relative positions of Moon and Sun in the sky, which determine the magnitude and variations of the total force of gravity and the lunisolar gravimetric tides, the effects of which are known to regulate several features of plant life (*42*), might provide a reliable cue of the approaching celestial event. In fact, gravitational forces on Earth are maximal when the Moon is simultaneously close or at its perigee (at the nearest point to the Earth) and syzygy (full or new moon), which is precisely the planetary configuration we observed on the day of the solar eclipse at the study site. This perigee-syzygy configuration, where the Moon passes exactly the same node at which its orbital plane intersects with the ecliptic Earth-Sun plane, characterising a certain Saros eclipse series (in our case, Saros 124) occurs every 18 years. Such a time-specific power surge in lunisolar gravimetric forces signalling an eclipse event could generate the anticipatory cue that older trees had experienced during similar events in the past (i.e. information stored as memory). Irrespective of the precise nature of the anticipatory cue, we found that the couple of older trees was in fact the most responsive in its anticipation. This response was expressed by the earlier rise in time asymmetry, a finding that is consistent with the process of transforming stored memories into information available in the present in open, dissipative systems (*43, 44*), such as trees.

Our understanding of collective behavioural phenomena, including the role that synchronisation plays in information processing and collective decision-making, has benefited enormously from the study of animal groups (Couzin 2018). Our results now show that synchronisation and collective memory are also an essential component of plant behaviour. In agreement with these results, the thermal state describing the tree and its environment (Eq. (S6) in Section S0.5 of Supplementary Materials) has built in (*45*) the Bayes’ probability rule by which, according to previous experiences, the action is estimated to be taken, among many possible ones, with higher (Bayes) probability in order to face an environmental event. We thus propose that despite the vast differences among organisms, the ability to transform information stored as collective memories into available information for collective decision-making, constitutes an adaptive mechanism shared across taxa, including plants. The occurrence of previously unexplored collective phenomena in plants suggests that the roles of synchronisation and collective memory in nature have been overlooked and underestimated. Our findings show that collective phenomena have emerged in both plant and animal groups, hinting at the intriguing possibility of a fundamental commonality in how patterns of behaviour propagate across different organisms.

## Acknowledgments

We acknowledge Alessandro Bernard, Paolo Ceretto, and Massimo Arvat, the Ente Parco Naturale di Paneveggio Pale di San Martino, Provincia Autonoma di Trento, the Administrative Director Fiorella Zortea, the entire Corpo Forestale at Stazione Forestale Demaniale di Paneveggio e Cadino, Girolamo Scarian, the entire workers’ squad, Piero Baldessari, Andrea Daprà, the *Genius loci* and *Salvanél*, Magnifica Comunità della Valle di Fiemme. Funding: main funding received from Zenit Arti Audiovisive, Torino (Italy); sponsorships received from OpenAzienda S.r.l., PrimoPrincipio Società Cooperativa, IGA Technology Services and Templeton World Charity Foundation. Author contributions: AC supervision, AC, PP, and SP experiments, AC, GV and MMD data analysis, GV theoretical aspects, AC and MG writing first draft, ALL discussion and final manuscript writing. Competing interests: the authors declare no competing interests. Data and materials availability: all data is available in the main text or in the supplementary materials.

