## Supplementary Materials for "Bioelectrical Synchronisation of Spruce Trees during a Solar Eclipse"

Andrew Adamatzky[2,3]

[1]Bioinspired Soft Robotics, Istituto Italiano di Tecnologia, Via Morego 30, 16163 Genova, Italy
[2]University of the West of England, Coldharbour Lane, Bristol, United Kingdom
[3]The Cyberforest Experiment, Paneveggio (TN), Italy
[4]The Biological Intelligence Lab, Southern Cross University, Lismore (NSW), Australia
[5]Openazienda S.r.l., Macomer (NU), Italy
[6]Dipartimento di Fisica "E.R. Caianiello", Università di Salerno, 84084 Fisciano (SA), Italy
[7]Departamento de Informática, Universidad Carlos III de Madrid, Leganés, Spain
[*]Alessandro Chiolerio;.

#### Supplementary materials

Contents:

General aspects

The Cybertree device

Higher-order complexity measures

Additional data

Some formal details of the theoretical modeling

#### S0.1 General aspects

For the collection of *xilematic* bioelectric potentials from one tree, ten stainless steel (AISI 316) threaded rods of 6 mm diameter were utilized. These rods were evenly spaced along the trunk, uncovered portion of roots and logs, with a separation distance of 50 cm. Each electrode pair was connected to a differential amplifier prior to recording the data.

For the collection of *phloematic* bioelectric potentials, two recording circles were positioned around the trunk of another tree. One circle was located 1 m above the ground level with a radial distance of $60°$, while the other was located 3 m above the ground level with a radial distance of $90°$. As with the *xilematic* potential collection, each electrode pair was connected to a differential amplifier prior to analog to digital conversion.

The same threaded rods were used to collect bioelectric signals from five logs. The first rod was inserted at the top of each log, and the second was positioned along one of the roots, 50 cm away from the first. Both rods were then connected to the differential amplifier. Signals were transmitted using the double-shielded ultra-low resistance INCA1050HPLC cable from MD Italy, designed for high-fidelity audio applications.

The electrodes were labelled with Greek letters $\alpha, \beta, \gamma, \delta, \epsilon, \eta, \zeta, \theta, \iota, \kappa$. The collection sites were labeled with Latin letters 'B', 'D', 'E', and 'G'. Collection sites 'B' nad 'D' represent a healthy, sun-exposed tree approximately 70 years old, site 'E' represents a portion of five dead logs in an area devastated by the 2018 Vaia storm, and site 'G' represents a healthy, shadowed tree approximately 20 years old. These collection sites were selected to showcase the varying ages and vitality levels that can be found in forests.

The Cybertree device consists of a device that can provide a remote reading of the biopotential from five differential electrode couples, as described in subsection S0.2. Further numerical techniques were utilized to extract complexity metrics from the biopotentials, as detailed in subection S0.3.

#### S0.2 The Cybertree device

##### S0.2.1 Hardware

CyberTree is a low-cost, low-power consumption, low-temperature resistant device designed for remote sensing the electrical signals of trees.

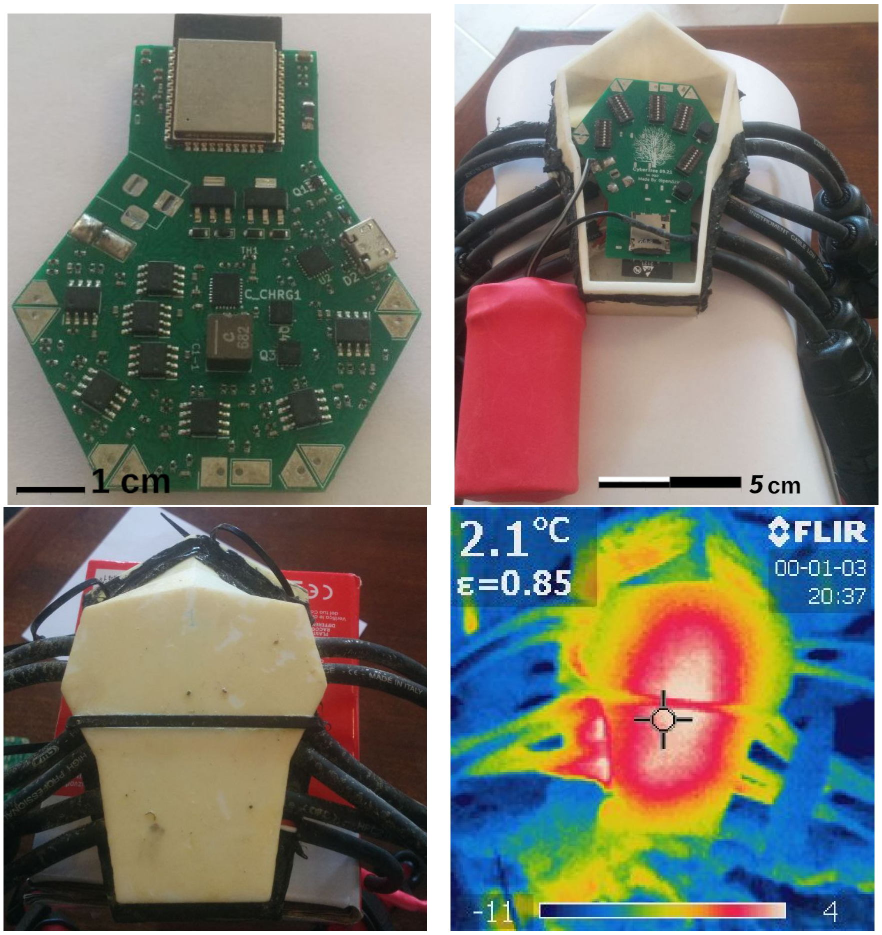


Figure S1: Different views of CyberTree device. From the top left, moving clockwise, images show the back view of the device, the device mounted into the case (with the red battery pack attached), the device in the final case, and an *in situ* thermal image of the working device.

It consists of:

- ESP32-WROOM32E (*1, 2*) MCU, memory: 16 MB

- 10 copper terminal (designed for easy-soldering the 5 electrodes couple)
- 5 operational amplifiers, each connected to an ADC channel.
- 5 signal translators (one for each signal channel)
- a 5 V power/programming USB interface
- microSD (TF card) storage
- battery charger circuit (for 2-cells, 6.4 V $LiFePo_4$ battery)
- 9 - 12 V power/charging interface

Table S1 shows the electric characteristic of the device.

| **Characteristic** | **Typical** | **Min, Max** | **Unit** |
|---|---|---|---|
| Commercial name | CyberTree | | |
| Manifacturer | OpenAzienda srl | | |
| N. of acquisition channels | 5 | | |
| Power Supply | 9 | 5, 12 | V |
| Operating Temperature | - | -40, 80 | °C |
| Sample rate (each channel) | 55 | | Hz |
| Max current (acquisition at 9 V) | 0.27 | 0.2, 3 | A |
| Max current (charging battery at 9 V) | 1.4 | 0.6, 3 | A |
| Dimension (board, L x H x W) | (60 x 79 x 10) mm | | |
| Dimension (case, L x H x W) | (80 x 130 x 43) mm | | |
| Weight (with batteries & case) | 250 g | | |

Table S1: General characteristic of CyberTree device

In our case CyberTree are placed in an outdoor environment, and exposed to meteorological events like rain, sun, and wind.

Internet connection is realized with a network star topology, with a wifi LTE router that works as a gateway for all the CyberTree devices.

Power is provided by a solar panel with batteries and it is used to power the LTE-wifi router and the CyberTree network.

**MCU** ESP32-WROOM32 ( (*1, 2*)) is used in CyberTree device as MCU, setting 5 channels of the first ADC (a 12-bits SAR ADC) for data acquisition and i2c connection with battery charging circuit (for charger circuit telemetry). It also provides the wifi remote connection and data storage via microSD interface and the PWM for the signal translator.

**Signal cable:** Electrodes are connected to the cybertree device using audio high-quality double-shielded cable for minimizing external noise. Cables are connected through IP68 joint to same-type cable, outgoing from CyberTree case, made by 3D-printed, temperature-resistant, plastic (figure S1).

**Data acquisition chain** for each channel is primarily composed of a low-noise, low-power audio op-amp in a summing and differential configuration for giving a differential signal with an offset (figure S2). ADC input signal is given by:

$$V_{sig} = 2 \cdot (S2 - S1) + Translator$$

The 2 factor is due to the opamp gain setting (via a resistor, see figure S2). CyberTree admits through parallel resistors, to manually switch the gain, changing the factor in the formula from 2 to 4 or 8. However, in this experiment, we use the standard configuration.

Our MCU can reach at ADC pin the maximum resolution of $\approx 250\mu V$ (at central voltage position).

Translators are obtained using a one-pole low-pass filter set for working at maximum PWM frequency (38 kHz), with 10-bit resolution. Signal processing, after the filter, continues with

| Attenuation (dB) | Suggested Range (mV) |
|---|---|
| 0 | 100 - 950 |
| 2.5 | 100 - 1250 |
| 6 | 150 - 1750 |
| 11 | 150 - 2450 |

Table S2: ADC characteristic of ESP32

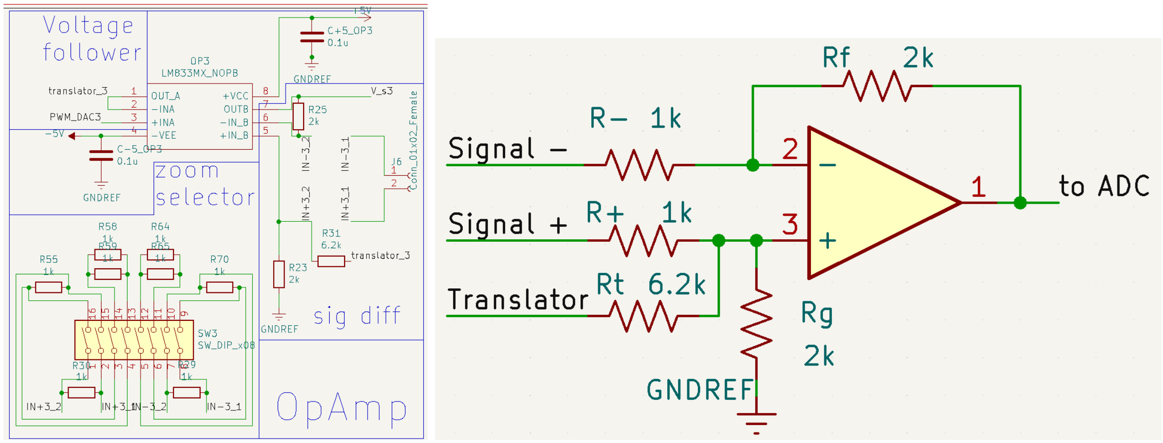


Figure S2: Schematics of op-amp configuration in CyberTree (left). The pwm2dac (input of the voltage follower) is the pwm line after the low-pass filter, `translator` is the signal after the voltage follower, input of the op-amp in summing configuration. An analog gain switch is given to set the signal gain to 2,4 or 8 (depending on how many parallel resistors are connected via dip-switches). Equivalent simplified schematics are shown at right (with gain = 2 for the input signal and translator scaled to max 1 V).

a voltage follower and input in the opamp as translator summing input. The CyberTree device also has an analog zoom switcher for the signal: changing the input equivalent resistance for the signal, is possible to switch between a gain of 2, 4, or 8.

In order to measure the negative value, a 5 V negative power supply (for op-amp dual powered mode) has been provided through micropump.

The final configuration with translators provides a possible acquisition range of [-1,1] V. In in this experiment we set up 0 db attenuation with a result in a range of observable signals of [-450,450] mV. With this configuration the signal is usually observed in the central part of the ADC range, reaching maximum resolution and best-condition measurement.

**ADC calibration.** ESP32 have per-device ADC voltage calibration parameter stored in esp32-specific internal registers ( (*1*)). Using a SIGLENT SDS 1202X-E oscilloscope, every device is tested in all the ranges for 0, 2.5, and 6 dB attenuations and compared with testing firmware, giving ADC count and the corresponding voltage by industry. These tests had confirmed the industry setting, with substantial agreement between the given voltage and experimental data from SIGLENT SDS 1202X-E oscilloscope.

**Translator calibration.** After achieving ADC calibration, a per-channel translator calibration is mandatory for obtaining data in a calibrated standard unit (mV) .
All the translators signal is scaled to fit the maximum value of 1 V.
CyberTree terminals are all connected to the same ground and we use the standard firmware for data acquisition, with a custom part for the translator. the duty cycle was set from 280/1024 to 1010/1024 (step: 10/1024) with oversampling of 1000 samples, giving 74 points for fitting results (each one with pwm setting and voltage from firmware). As validation, each signal is also verified using the oscilloscope. For example, a translator-voltage calibration fit for all channels in a single CyberTree device is shown in figure S3.

All the CybertTree used are tested with the same procedure, giving the linear coefficients for the firmware. Our transation level is given by

$$V(mV) = (0.76 \mp 0.01) \cdot (x) + (35 \mp 3)$$

when x is the pwm duty cycle set (from 0 to 1024) and the error is obtained taking into account all the differences between channels and devices. During this calibration, no correlation

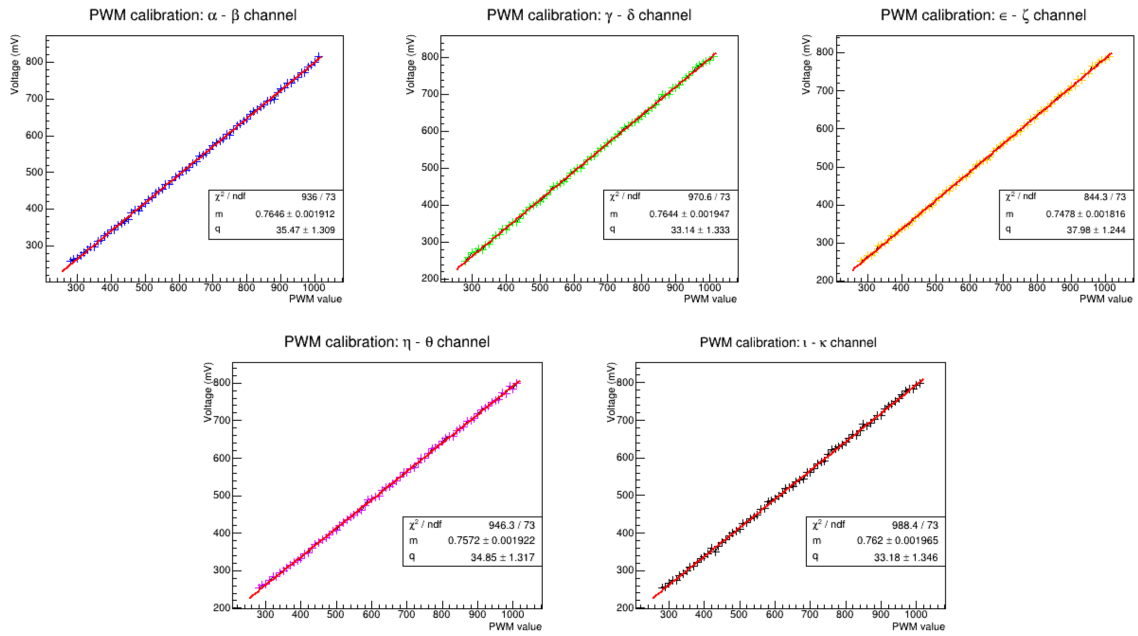


Figure S3: Example of calibration for PWM values of one CyberTree device. Similar calibrations were made for all the devices, for testing the reproducibility and checking the translator features for a single channel in each device. This test demonstrates the same calibration values for all the devices and all channels.

between the specific channels and experimental bias has been detected.

**Data storage:** Since the MCU has limited memory, CybreTree integrates a microSD card interface for storing data. In our application, data are stored before sending it to the cloud (for energy saving, we send data every 5 minutes).

**Energy management:** The CyberTree has a multiple power supply implementation, with a 5V supply channel (USB interface) for operating without a battery, and an external supply implementation connected to the battery charger. This circuit provides a power path feature (when an external supply is provided, the battery is in charge mode and the power path provides the supply for the device). The device provides also some information by remote request: charging state, battery voltage, input current, and output current.

Used batteries are 2-cells, 6.4 V, 3200 mAh $LiFePO_4$ batteries.

**Energy system**

In our installation, the power supply is provided by the system shown in figure S4, CyberTree devices work and charge on-device batteries with a central supply (during solar-powered time). The central power station can detach the CyberTree Line, keeping the power line only on the router when CyberTree devices work battery-powered.

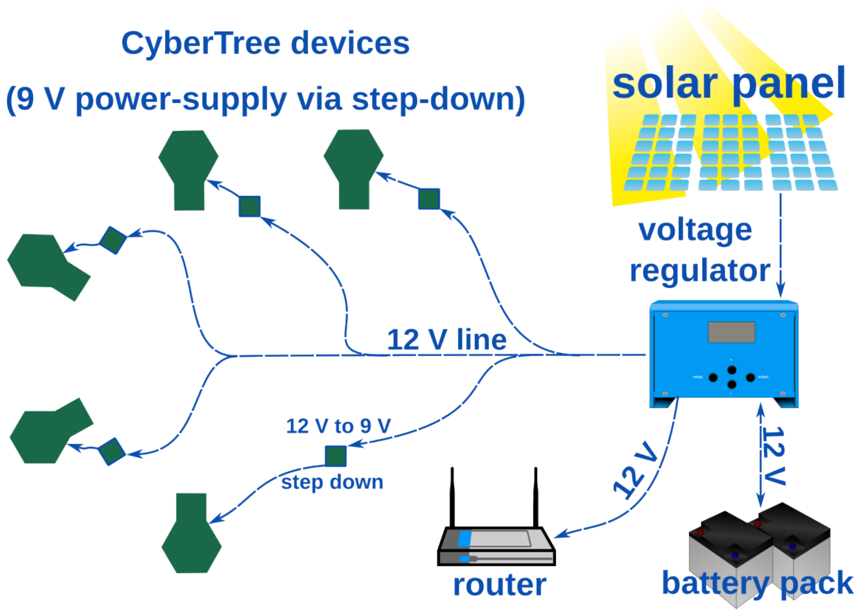


Figure S4: Power supply scheme for this experiment: a two solar panel system (max power 130 W + 50 W) is connected to a voltage regulator for power supply and battery charging, battery pack (12V 180 Ah) provides energy storage. The router is directly connected to the voltage regulator. 12V powered line with terminal step down to 9V are used for easy design and optimal voltage supply for CyberTree devices

For energy sizing and power estimation, the EU JRC Photovoltaic Geographical Information

System with PVGIS 5.2 ( (*3*)) was used.

The input parameters for this eximation are shown in table S3 And giving the outputs:

| **Parameter** | **Value (unit)** |
|---|---|
| System Type | Off-grid |
| Latitude | 46.316 |
| Longitude | 11.744 |
| Database used | PVGIS-SARAH2 |
| PV installed | 180 (Wp) |
| Slope | 30 (°) |
| Azimuth | -70 (°) |
| Battery capacity | 2160 (Wh) |
| Batt. Cutoff limit | 20 (%) |
| Consumption per day | 288 (Wh) |

Table S3: Input parameter for estimating power statistics of 5 CyberTree system with the outdoor router for remote communication

- **Days with full battery:** 52.16 %
- **Days with empty battery:** 17.07 %
- **Average energy not captured:** 331.79 Wh
- **Average energy missing:** 119.16 Wh

These settings result in energy independence with 24h activity from March to October. From November to February the system was normally shut down and powered on remotely (by turning on some remote-controlled relays).

##### S0.2.2 Firmware

The Firmware of CyberTree is realized with microPython language. It supports object-oriented programming and async input-output methods. Micropython port for ESP32 use FreeRTOS

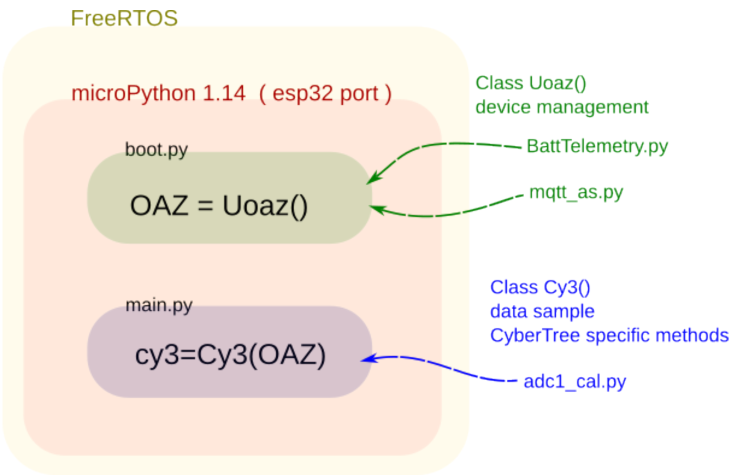


Figure S5: CyberTree firmware diagram, main classes and general architecture for micropython port of ESP-32. Uoaz class init device in `boot.py` and Cy3 implement specific device method in `main.py`, imported classes are shown at the right part of the figure.

(figure S5), at the time of firmware development, the micropython version for ESP32 was 1.14 ( *(4)*). Methods are divided in two main classes:

- **Uoaz**: including all the methods for general device management (WiFi and Cloud connectivity, remote management, configuration updates, remote firmware upgrade). This class is created in the `boot.py` and is run once at startup.
- **Cy3**: specific implementation of CyberTree device (Data acquisition firmware, remote message format, timing for data-sending). This class is created in `main.py` and implements the main loop.

The other 3 classes provide specific implementation:

- `BattTelemetry.py` implementation of Battery telemetry
- `mqtt_as.py` asynchronous implementation of mqtt client-side protocol
- `adc1_cal.py` calibrated conversion from ADC count to real voltage

The first two classes are integrated into Uoaz class. `adc1_cal.py` is integrated in Cy3. CyberTree design takes into account the telemetry and remote management of the device, used in our experiment. It requires an internet connection. Uoaz class implements a fallback mode in case of an unavailable internet connection, offering a local web interface for in situ configuration.

The fallback mode can be used to set up WiFi credentials and enable the device for a first connection to the internet. After it, Uoaz class is responsible for setup remote management features, including remote configuration and firmware upgrades.

##### The Cy3 class and the data sample implementation

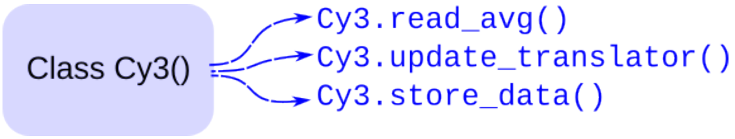


Figure S6: Cy3 Class and relevant methods used for data sampling, translator management, and telemetry during this work.

Cy3 class implements specific CyberTree methods and it is responsible for data sampling (as hown in figure S6). Routines are designed in order to ensure the two-parameter of sample precision: time and voltage.

Time precision is implemented in two ways: NTP protocol for common device synchronization, a time counter, and a deadline system to ensure the sampling cycle stays in the defined period of 1 second (sample rate 1 Hz). Some parameters are defined by configuration and have been tuned in the laboratory.

Voltage precision is improved with different strategies: oversampling, translation and zoom of the signal.

**Cy3 initialization:** Initialization routine primarily set all data acquisition parameters, for doing it, Cy3 class:

1. initializes one adc1_cal class for each channel by setting the ADC pin.
2. setup all ADC resolutions to 12 bits
3. setup ADC attenuation, getting values from configuration file
4. initialize translator's PWM with a starting duty-cycle of 512/1024.
5. initialize samples vector to zero

**Configuration** is stored in json format in the root of esp32 flash memory. Json is created via a web interface on the cloud server and can be modified remotely from the cloud.

Sampling parameter for the used configuration are:

- CY3_ATTEN = 0 (Db) : ADC attenuation
- CY3_LINE2SEND = 300 : Line of data for each send
- CY3_SAMPLEPERIOD = 20 (ms) : period for each data acquisition for oversampling
- CY3_SAMPLEPERLINE = 55 : oversampling factor

Other configuration parameters are used for telemetry and remote management.

**The sampling method: `Cy3.read_avg():`** ADCs are configured for the maximum resolution of ESP32 (12 bit). An acquisition is perfomed with an average from a set of n (CY3_SAMPLEPERLINE, 55) for each second. Between each single sample esp32 machine wait for a period of CY3_SAMPLEPERIOD (20) ms.

Those values have been defined in a fine tuning of device. In order to ensure time precision, overhead used for composing and sending message is estimated in 80 ms. Before of each acquisition the timestamp is recorded in a variable that is incremented by 920 ms and rappresent a deadline for current acquisition. The acquisition is perfomed CY3_SAMPLEPERLINE times with a sample-period between each. In case of any delay the acquisition is completed with a minor number of sample in order to ensure the time precision. This behaviour is achieved in cycle that loop for average estimation: a break is triggered in case of deadline or after CY3_SAMPLEPERLINE sampling. In order to optimize use of memory, averages are perfomed with in place summing of values. At the deadline, sum is divided by the effective number of samples recorder in that second.

The oversampling implemented improves acquisition by $\sim 2.9$ bits. We can add this value to the 12-bit of ADC with a final resolution of 14.9 bits for the range observed by the ADC (950 mV at 0 dB attenuation). It results in a precision of $\sim 31\mu V$ of ADC input signal (in this work opamp amplify the signal by gain = 2, giving $\sim 15.5\mu V$ )

**The translator adjustment method : `Cy3.update_translator()`**

In order to have the signal at the central window of ADC admitted values, and include both positive and negative voltage in the range of measurement, a translator is realized with a combination of PWM and a filter circuit (as described in paragraph S0.2.1 ). This method is responsible for setup the value of the duty cycle of each PWM, and causing the effective value of the signal translator paired with each ADC. The value is updated each time the signal moves to the upper or lower part of ADC. The method verifies the value of the last sample and the value of the duty cycle. In case the duty cycle is greater than an upper threshold and the signal measured by ADC is greater than 800mV the duty cycle is decreased. Otherwise, in case is lower than the lower threshold and the signal measured by ADC is lower than 250mV the duty cycle is increased.

In this way, we can concentrate the ADC precision in a part of the signal and reach a better final precision in acquisition.

The value obtained with a translator and its relation with PWM duty cycle has been verified and calibrated in the laboratory (see paragraph S0.2.1)

##### The store method: `Cy3.store_data()`

The store_data method is used to perform acquisition for CY3_LINE2SEND (300) times and send the data with mqtt, or, in case of configuration with an SD memory, to store data in an sd card. In the actual configuration, firmware is set to work with telemetry and send data over the internet with mqtt protocol. For this purpose, each device has a configured MQTTCLIENT-NAME, and a dedicated MQTTDATATOPIC in a remote server.

`Cy3.store_data()` method is designed to realize a set of acquisitions calling the function `Cy3.read_avg()` CY3_LINE2SEND times. The variable is used to fine-tune the device and avoid a continuous transmission for each acquisition, in order to reduce overhead, settings for this work define a data transmission every 5 minutes.

This function is also responsible to call the `Cy3.update_translator()` method and adjust the translation of the signal.

##### Uoaz class and the device management implementation

Device management is a critical design factor in order to run experiments in an outdoor environment, like in our experiment. CyberTree design takes into account the following features:

- remote diagnostic (including battery telemetry)
- remote configuration update
- remote firmware upgrade
- remote reboot

Class Uoaz is responsible for startup and ensuring a correct clock setup with Network Time Protocol (NTP, (*5*)) in order to achieve for each device a common synchronization.
The data sent to the server have the same time reference and each sample, is expressed in second (Epoch Unix Timestamp format).
Each CyberTree device has an initialization that checks network availability to a specific WiFi network defined in the configuration.
Once connected, the device authenticates itself to the remote server: each device has a unique identifier that is generated with a random Universal Unique Identifier (UUID-v4, (*6*)). This identifier is used for authentication to a remote server with POST to a server endpoint, MAC address of the esp wifi interface is included in the POST. The server accepts only devices that match with the known pair of UUID and MAC.
Authenticated devices receive a JSON with the last configuration setup defined in the web interface of the server.
The configuration includes a set of sampling parameters plus a set of device management parameters.
Mqtt ( (*7*)) is used for two purposes: data-telemetry (Cy3 class) and device management (Uoaz

class).

Configuration includes the setup of two MQTT topics dedicated to remote management: the MQTTCTRLTOPICsub and the MQTTCTRLTOPICpub. The first is configured for subscription in order to receive the commands from the remote server, the second is used to publish responses for commands.

Remote management are provided different features:

- device announcement
- battery telemetry
- reboot
- configuration update
- firmware upgrade

An announcement of the device is performed at each device boot and is used to check the actual status of the CyberTree network.

Battery telemetry is performed when the remote administration of the device sends the command via the web interface.

In order to perform an upgrade of the firmware reboot is required, causing the CyberTree to check for new firmware at startup.

#### S0.3 Complexity measures

In various demanding signal processing systems and applications, relying solely on second-order statistics such as the mean, variance, or correlation proves inadequate when the data deviates from a Gaussian distribution and the adaptive system exhibits nonlinearity. In such scenarios, accurately representing linear/nonlinear adaptive signal processing systems requires incorporating higher-order statistics. Furthermore, the concept of information theory indicates that the uncertainty level influences the information value of data. Data corresponding to highly probable events carries limited information, while data associated with less probable events carry more information. Higher-order statistics, through metrics such as entropy, Simpson diversity, expressiveness, and Lempel-Ziv complexity, assist in reducing uncertainty (*8–11*).

In this study, we compute the following information-theoretic complexity measures to characterise the spatiotemporal activity patterns in raw data within a 10-minute time window.

1. The Shannon entropy ($\mathcal{I}_1$) serves as a measure of uncertainty for discrete random variables. Given a random variable $s$ with $n$ elements, $s = \{s_1, s_2, \cdots, s_n\}$, and its probability distribution $p(s) = \{p(s_1), p(s_2), \cdots, p(s_n)\}$, the mathematical expression for Shannon entropy is given by Equation S1.

$$\mathcal{I}_1 = -\sum_{i=1}^{n} p(s_i) \log\left(p(s_i)\right). \tag{S1}$$

2. The Rényi entropy ($\mathcal{I}_2$) is a measure of diversity and plays a fundamental role in statistics (*12*). The mathematical formulation for this entropy, with a fixed parameter $q$ set to 2, is provided in Equation S2.

$$\mathcal{I}_2 = \frac{1}{1-q}\left(\ln\left(\sum_{i=1}^{n} p(s_i)^q\right)\right) \tag{S2}$$

3. The Tsallis entropy ($\mathcal{I}_3$), which generalises the standard Boltzmann-Gibbs entropy, represents a non-extensive entropy measure (*13*). The mathematical expression for Tsallis entropy, with $q$ and $k$ representing the degree of non-extensivity and a positive constant, respectively, is given in Equation S3. In our empirical study, we adopted the setting of $q = 2$ and $k = 1$.

$$\mathcal{I}_3 = \frac{k}{1-q}\left(1 - \sum_{i=1}^{n} p(s_i)^q\right) \tag{S3}$$

4. The Space filling ($\mathcal{I}_4$) is defined as the ratio of non-zero elements in the signal $s$ to the total length of the signal.

5. The Expressiveness ($\mathcal{I}_5$) is the ratio of Shannon entropy ($\mathcal{I}_1$) to Space filling ($\mathcal{I}_4$). This metric provides insight into the "economy of diversity."

6. The Diversity index ($\mathcal{I}_6$) quantitatively evaluates the number of unique activities of interest in the acquired signal, considering aspects such as richness, divergence, and evenness. Its mathematical formulation, with the parameter $q$ set to 3, is given by Equation S4.

$$\mathcal{I}_6 = \frac{1}{\sqrt[q-1]{\sum_{i=1}^{n} p(s_i)p(s_i)^{q-1}}} = \left(\sum_{i=1}^{n} p(s_i)^q\right)^{1/1-q} \tag{S4}$$

7. The Simpson Diversity ($\mathcal{I}_7$) measures the concentration of individuals classified into types and is calculated as $\mathcal{I}_7 = \sum_{i=1}^{n} p(s_i)^2$. The range of $\mathcal{I}_7$ is between 0 and 1, with 1 indicating infinite diversity and 0 representing no diversity.

8. The Lempel–Ziv complexity ($\mathcal{I}_8$) is employed as a measure of temporal signal diversity and focuses on compressibility. The Kolmogorov complexity algorithm (*14*) was used to calculate this metric, which provides a scalar measurement for estimating the bandwidth of random processes and quantifying the harmonic variability in quasi-periodic signals.

9. The Perturbation Complexity Index (PCI) ($\mathcal{I}_9$) normalises the Lempel–Ziv complexity of the spatiotemporal pattern of the signal with respect to its Shannon entropy ($\mathcal{I}_1$).

10. The Kolmogorov complexity ($\mathcal{I}_{10}$) assesses the information content of a signal by quantifying the minimum length of a binary code required to describe it with an optimal encoding. The mathematical expression for Kolmogorov complexity is presented in Equation S5.

$$\mathcal{I}_{10} = \min\{|\mathcal{P}| \mid \mathcal{P} \in 0, 1^*, U(\mathcal{P}) = s\} \tag{S5}$$

where $U(\mathcal{P})$ is the output of the binary program $\mathcal{P}$ when executed on a fixed reference universal Turing machine and $|\mathcal{P}|$ is the length of the binary program $\mathcal{P}$.

11. The Fractal dimension ($\mathcal{I}_{11}$) measures the self-similarity of a signal across multiple scales. We use the Higuchi method (*15*) in this study to calculate the fractal dimension of a signal, as represented by $\mathcal{I}_{11} = \frac{\log(n)}{\log(1/d)}$. Here, the parameter $d$ denotes the step size used in signal sampling, and $\log(\cdot)$ represents the natural logarithm.

#### S0.4 Additional data

Data visualisation: Savitzki-Golay $1^{st}$ order smoothing over 51 points is superimposed on each recording of Figure S7. The data chunk spans 120.000 s and features five differential channels (curves) plus the "ON time" of the device (first curve from the top of panel A), information useful to correlate with spikes that might occur during eventual restart.

Additional measurements are related to sites B, E and F: the latter features phloematic differential channels and is a tree older than 70 years. In Figure S7 we show an exemplar recording of 120.000 s, from all of the differential couples, in panel a). The dashed red arrows locate the starting and ending point of the solar eclipse, showing that two phenomena visible trends occurred: a first, slow change of the initial biopotential level by approximately $300\%$ happening during the first 50.000 s. This is evident from the bottom circle electrodes (pairs $\epsilon - \eta$, $\theta - \zeta$, $\iota - \kappa$), where the couple facing the sun and oriented southwards ($\epsilon - \eta$) experiences an increase of the biopotential, while the other two couples experience a reduction. See also the noise level that is higher in the first case. This trend is not found in the couples belonging to the top circle ($\alpha - \beta$ and $\gamma - \delta$). A second trend, directly associated with the geometric occultation of the Sun and enclosed in the red arrows, corresponds to a reduction of the biopotential, and is sharp and evident, particularly for the couple $\gamma - \delta$. This same trend can also be found in the three couples of the bottom circle, taking place over a slightly broader timescale, but yet visible. The last part of the measurements, after the ending of the eclipse until 120.000 seconds, features a renormalization of the biopotential levels, that approach their initial values, on all of the couples. The magnified portion of the biopotentials of the couple $\alpha - \beta$ shown in panel b) gives also an idea of the variation in the noise floor, before and after the eclipse. To better put in evidence this aspect, a Fast Fourier Transform (FFT) analysis was performed, comparing a chunk of 40.000 seconds before the eclipse (blue curve) with a chunk taken after the eclipse. An important reduction of the noise power is seen, particularly at low frequencies, corresponding to the suppression of oscillations with periods between 680 s (1.47 mHz) and 111 s (9 mHz). Also, the Short Time Fourier Transform (STFT) map shown in panel d) gives a qualitative view of the effects of the solar eclipse on the biopotential signals.

For what concerns the E site, it features five dead logs, with xilematic potential recordings, while the B site features a tree of approximately 70 years of age, with xilematic potential recordings. We see very little variations in their bioelectric response (see Figure S8). Only a deep analysis, using FFT tools, tells us that there is actually a difference in the biopotential spectrum before and after the eclipse, but this difference goes in an opposite direction, with respect to the previous two cases of the phloematic recordings. We see that the oscillation frequency increases after the cosmologic event. Being the xilematic electrodes connected more to the inner physiology of the plant, and to the movement of fluids inside the trunks triggered by capillarity and water pressure in the soil, we deduce that a higher frequency in the oscillation of liquids is generated. Oscillating masses reduce their fluctuation periods when the mass increases, therefore it is likely that the effective mass has increased.

#### S0.5 Some formal details of the theoretical modeling

Since trees are open, dissipative systems, the QFT canonical formalism requires (*16–19*) that also the environment has to be included in the description. This can be obtained by using

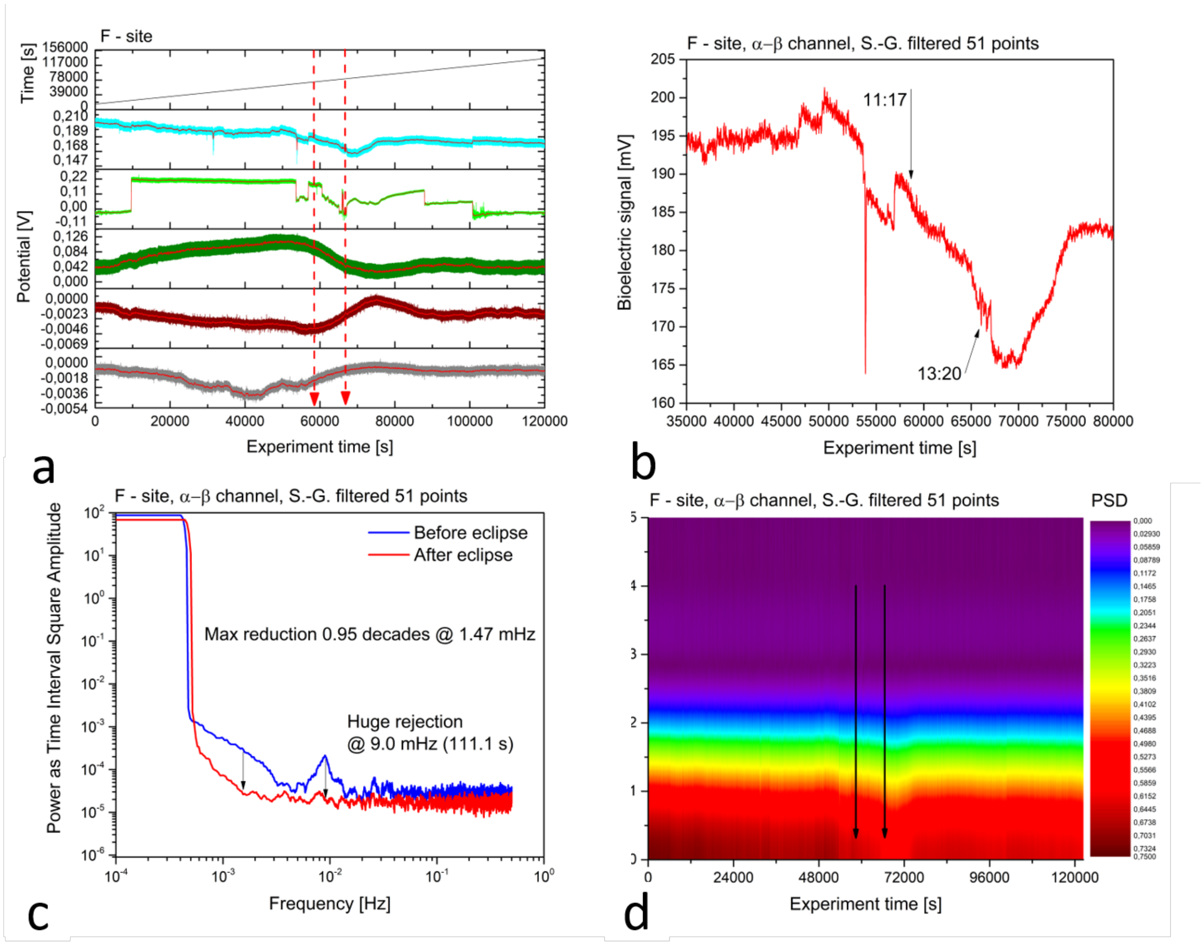


Figure S7: a) Raw measurements collected from the five differential channels, phloematic signals. b) Enlarged view of the $\alpha - \beta$ differential channel with solar eclipse timing indicated. c) FFT analysis of $\alpha - \beta$ differential channel. d) STFT analysis.

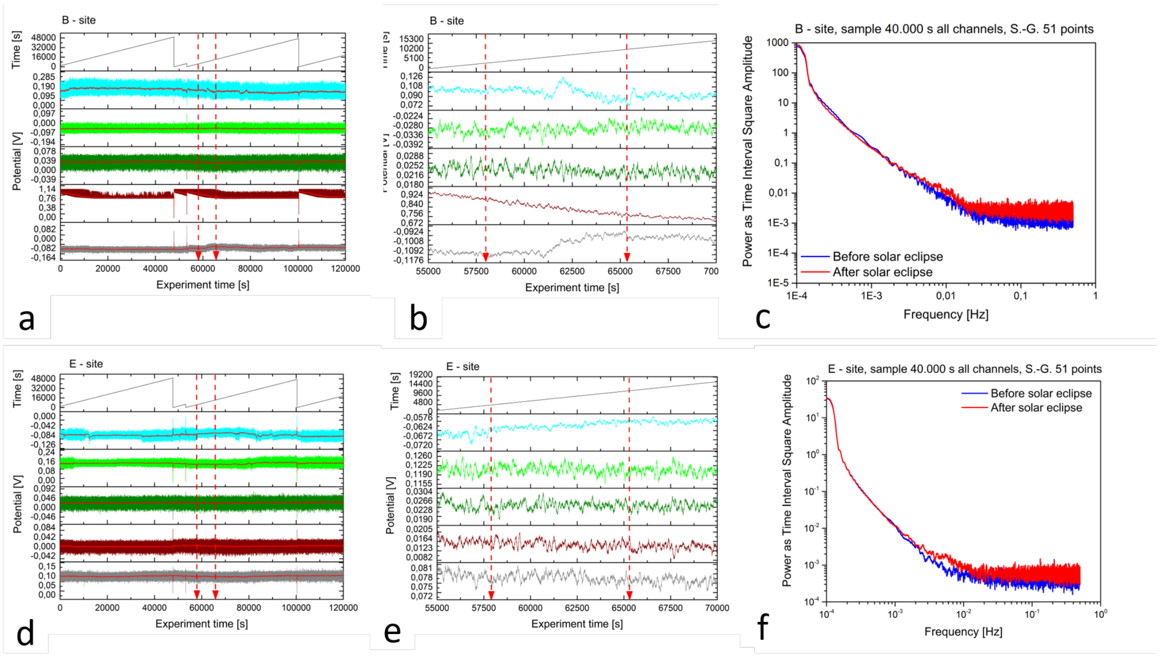


Figure S8: Bioelectric potentials recorded from sites B and E during the solar eclipse. a) raw measurements The red arrows indicate the starting (11:17) and the ending (13:20) of the solar eclipse. b) Enlarged view of the differential channels. c) FFT analysis. d) raw measurements collected from the five differential channels connected to the logs in site E. e) Enlarged view of the differential channels. f) FFT analysis.

the well known formalism of finite temperature QFT, also called the Thermo Field Dynamics (TFD) formalism. For completeness, we briefly summarize here few essential details of such a formalism which are useful to our theoretical modeling. A complete account of TFD in QFT can be found in (*16, 17*), see also (*18*).

The QFT state describing the system and its environment is (*19–21*)

$$|0(\theta(\beta))\rangle = \prod_{\mathbf{k}} \frac{1}{\cosh\theta_k(\beta)} \exp\left(\tanh\theta_k(\beta)\, a^\dagger_{\mathbf{k}}\tilde{a}^\dagger_{\mathbf{k}}\right)|0\rangle. \tag{S6}$$

It is normalized to 1, $\langle 0(\theta(\beta))|0(\theta(\beta))\rangle = 1,\ \forall\theta(\beta),\ \forall\beta,\ \forall t$. For notational simplicity, the time-dependence of $\beta$ is not shown: $\beta \equiv \beta(t) = 1/k_B T(t)$. The state $|0(\theta(\beta))\rangle$ is the $SU(1,1)$ generalised coherent state (*22*) at finite temperature $T$. It is an entangled and squeezed state. The squeezing parameter $\theta_k(\beta)$ (*18, 23*) is related to the fractal dimension in the isomorphism between fractal self-similarity and deformed coherent states (*24,25*). ${a^\dagger}_{\mathbf{k}}$ and $a_{\mathbf{k}}$ are the creation and annihilation operators, respectively, of the NG long range correlation field introduced in the main manuscript. $\tilde{a}^\dagger_{\mathbf{k}}$ and $\tilde{a}_{\mathbf{k}}$ are the creation and annihilation operators representing the environment (the thermal bath) (*16–19*).

In QFT with SBS the ordered patterns dynamically generated by the long range correlations are characterised by the "order parameter" (in our case the polarization density $P(x,t)$), which is a classical field, meaning that it is not affected by quantum fluctuations, although it is of quantum origin. The macroscopic (classical) description (associated with the behavior of the classical order parameter field) is thus dynamically obtained from the microscopic (quantum) one.

Let $|0\rangle$ denote the state without condensed couples $(a_{\mathbf{k}}, \tilde{a}_{\mathbf{k}})$. States at different temperatures, $\beta \neq \beta'$, are orthogonal states in the limit of infinite volume $V \to \infty$ (*16–18*):

$$\langle 0|0(\theta(\beta))\rangle \to 0, \quad \text{and} \quad \langle 0(\theta(\beta'))|0(\theta(\beta))\rangle \to 0, \quad \text{for } V \to \infty \quad \forall\beta \neq \beta', \tag{S7}$$

which means that they belong to unitarily inequivalent representations of the canonical commutation relations. Thus, the change $\beta \to \beta'$ leads to different dynamical regimes, $|0(\theta(\beta))\rangle \to |0(\theta(\beta'))\rangle$ (*16–18*).

The covariance $cov(N_a, N_{\tilde{a}}) \equiv \langle N_a N_{\tilde{a}} \rangle - \langle N_a \rangle \langle N_{\tilde{a}} \rangle$ provides a measure of the degree of entanglement (the symbol $\langle ** \rangle$ denotes expectation values in $|0(\theta(\beta))\rangle$; for simplicity, the subscript $\mathbf{k}$ has been omitted). One gets $cov(N_a, N_{\tilde{a}}) = (1/4)\sinh^2\theta \neq 0$ for $\theta \neq 0$. Instead, $cov(N_a, N_{\tilde{a}}) = 0$, $\forall\ \theta$, for non-correlated modes since $\langle N_a N_{\tilde{a}} \rangle = \langle N_a \rangle \langle N_{\tilde{a}} \rangle$ in that case (*20, 23, 26–28*).

One defines the entropy $S_a$ and $S_{\tilde{a}}$ (*16–18*), and

$$\langle 0(\theta(\beta))|S_a|0(\theta(\beta))\rangle = \sum_{n=0}^{+\infty} W_n \, \log W_n \, , \tag{S8}$$

$$W_n = \prod_{\mathbf{k}} \frac{\sinh^{2n_{\mathbf{k}}} \sigma_k}{\cosh^{2(n_{\mathbf{k}}+1)} \theta_k} \, , \tag{S9}$$

where $n$ denotes the set $\{n_{\mathbf{k}}\}$, $0 < W_n < 1$ and $\sum_{n=0}^{+\infty} W_n = 1$ (and similarly for $S_{\tilde{a}}$). Note that $dS_a = dS_{\tilde{a}}$ is implied by the fact that the difference $S_a - S_{\tilde{a}}$ is a constant of motion, expressing the dissipative coupling system-environment. Also note that higher is the entanglement (the covariance going as $(1/4)\sinh^2\theta$), lower is the entropy (cf. Eq (S9)). The free energy is

$$\mathcal{F}_a \equiv \langle 0(\theta(\beta))|\Big(H_a - \frac{1}{\beta} S_a\Big)|0(\theta(\beta))\rangle \, , \tag{S10}$$

where $U_a \equiv H_a = \sum_{\mathbf{k}} \hbar\omega_k a_{\mathbf{k}}^{\dagger} a_{\mathbf{k}}$ (and similar expression for $\tilde{a}$ modes). In the (quasi-)stationary case, the minimization condition $\partial \mathcal{F}_a / \partial \theta_k = 0$ , $\forall\ k$ , gives the Bose–Einstein distribution

$$\mathcal{N}_{a_{\mathbf{k}}}(t) = \sinh^2 \theta_k(\beta(t)) = \frac{1}{\mathrm{e}^{\beta(t)\hbar\omega_k} - 1} \, , \tag{S11}$$

for the $a_{\mathbf{k}}$ modes condensed in $|0(\theta(\beta))\rangle$. $\mathcal{N}_{a_{\mathbf{k}}}(t) \equiv \mathcal{N}_{a_{\mathbf{k}}}(\theta(\beta(t)))$ is the number of $a_{\mathbf{k}}$'s. We also obtain Eq. (3) in the main manuscript showing that to a decrease of entropy (i.e. increase of ordering) corresponds a decrease in the kinetic energy.

Time evolution of the state $|0(\theta(\beta))\rangle$ is controlled by entropy:

$$\frac{\partial}{\partial t}|0(\theta(\beta))\rangle = -\left(\frac{1}{2}\frac{\partial S_a}{\partial t}\right)|0(\theta(\beta))\rangle, \tag{S12}$$

expressing the irreversibility of time evolution (*the arrow of time*) (and similarly for $S_{\tilde{a}}$).

For small fluctuations $\delta\theta_{\mathbf{k}} = \theta'_{\mathbf{k}} - \theta_{\mathbf{k}}$ ($\theta_{\mathbf{k}} \leftrightarrow \theta'_{\mathbf{k}}$), and negligible $\partial\delta\theta_{\mathbf{k}}/\partial t$, around the condition $d\mathcal{F}_a = 0$, the entropy variation is

$$\Delta \sum_{\mathbf{k}} \hbar\,\omega_k\,\dot{\mathcal{N}}_{a_{\mathbf{k}}}(t)dt = \sum_{\mathbf{k}} 2\dot{\theta}_{\mathbf{k}}(t)\cosh(2\theta_{\mathbf{k}}(t))\delta\theta_{\mathbf{k}}dt = \frac{1}{\beta}(dS'_a - dS_a), \tag{S13}$$

which is related to the Kolmogorov entropy in nonlinear dynamical systems (*29*).

Notice that the state $|0(\theta(\beta))\rangle$ has built in the Bayes' probability rule (*30*). In fact, the probability to find $n_{\mathbf{k}}$ particles $a_{\mathbf{k}}$ given the presence of $\tilde{n}_{\mathbf{k}}$ particles $\tilde{a}_{\mathbf{k}}$ in the state, conditioned by the value $\theta_k$ satisfying the Bose-Einstein distribution function Eq. (S11), is

$$P(n_{\mathbf{k}}|\tilde{n}_{\mathbf{k}}) = \frac{\tanh^{2n_{\mathbf{k}}}\theta_k}{\cosh^2\theta_k} \quad \forall \mathbf{k}\,. \tag{S14}$$

(cf. Eq. (S9)). The same is also true for $\tilde{a}_{\mathbf{k}}$, since $a_{\mathbf{k}}$ and $\tilde{a}_{\mathbf{k}}$ come in pairs $\forall\mathbf{k}$. Thus, $P(n_{\mathbf{k}}|\tilde{n}_{\mathbf{k}}) \equiv P(\tilde{n}_{\mathbf{k}}|n_{\mathbf{k}})$. The probability to find $n_{\mathbf{k}} = \tilde{n}_{\mathbf{k}}$ particles in $|0(\theta(\beta)\rangle$ is $P(n_{\mathbf{k}}) = P(\tilde{n}_{\mathbf{k}})$, $\forall\mathbf{k}$. Then,

$$P(n_{\mathbf{k}}|\tilde{n}_{\mathbf{k}}) = \frac{P(\tilde{n}_{\mathbf{k}}|n_{\mathbf{k}})P(n_{\mathbf{k}})}{P(\tilde{n}_{\mathbf{k}})}, \tag{S15}$$

i.e. the Bayes' relation.

A final remark is on the asymmetry in the behavior of trees shown by Eq. (2) of the main manuscript. In full generality, suppose that among the many differences between trees there is one property which accounts at the macroscopic level of the complex molecular dynamics of the tree and that we assume might take two values, say "+" or "−". For example, let $(+)$ and $(-)$ be associated to molecular dynamical features of tree older than 80 years, and younger than that, respectively. The global energy of the molecular states of the trees has to be a scalar quantity, independent of (invariant under) rotations in the space of the two component vectors (doublets) $\mathbf{d}_i = (+ \quad -)_i^T$, with $i = 1, 2, ...$ running over the number of the trees in the forest, i.e. it should depend on scalar terms like $g \ \mathbf{d}_i \cdot \mathbf{d}_j$, for any $i \neq j$, $i, j = 1, 2, ..$; $g$ denotes the coupling constant and $\mathbf{d}_i \cdot \mathbf{d}_j$ the scalar product of $\mathbf{d}_i$ and $\mathbf{d}_j$. The lowest energy contribution from such terms comes from the component $(1/\sqrt{2})[(+)(-) - (-)(+)]$, corresponding to the total $\mathbf{d}_{ij} = \mathbf{d}_i + \mathbf{d}_j = 0$, and is indeed anti-symmetric under exchange $(+) \leftrightarrow (-)$. It is in fact $\mathbf{d}_{ij}^2 = (\mathbf{d}_i + \mathbf{d}_j)^2 = \mathbf{d}_i^2 + \mathbf{d}_j^2 + 2\mathbf{d}_i \cdot \mathbf{d}_j$, and considering that $\mathbf{d}_i^2 = (1/2)(1/2 + 1)$ and same for $\mathbf{d}_j^2$, we have $\mathbf{d}_i \cdot \mathbf{d}_j = -3/4$, provided $\mathbf{d}_{ij}^2 = 0$. In contrast, the energy contribution is higher for the symmetric case, $(1/\sqrt{2})[(+)(-) + (-)(+)]$, the eigenvalue of the total $\mathbf{d}_{ij}^2$ being now $2$ and thus $\mathbf{d}_i \cdot \mathbf{d}_j = (1/2)(2 - 3/2) = +1/4$. The conclusion is that the anti-symmetric case is favored since it corresponds to lower energy: in the forest there is no symmetry under exchange among $(+)$-trees and $(-)$-trees and Eq. (2) holds when one $(+)$- and one $(-)$-tree are involved.

### References


1. Espressif, Esp32_technical_reference_manual_en.pdf.

2. Espressif, Esp32-wroom-32_datasheet_en.pdf.

3. European Commission, JRC Photovoltaic Geographical Information System (PVGIS).

4. Damien P. George, Paul Sokolovsky, MicroPython 1.14.

5. D. Mills, J. Martin, J. Burbank, W. Kasch, Network Time Protocol Version 4: Protocol and Algorithms Specification.

6. P. Leach, M. Mealling, R. Salz, A Universally Unique IDentifier (UUID) URN Namespace.

7. Andrew Banks, Rahul Gupta, MQTT Version 3.1.1.

8. M. M. Dehshibi, A. Adamatzky, *Biosystems* **203**, 104373 (2021).

9. M. M. Dehshibi, *et al.*, *ACS Biomaterials Science & Engineering* **7**, 3718 (2021).

10. A. Chiolerio, M. M. Dehshibi, D. Manfredi, A. Adamatzky, *Biosystems* **218**, 104691 (2022).

11. M. M. Dehshibi, T. Olugbade, F. Diaz-de Maria, N. Bianchi-Berthouze, A. Tajadura-Jiménez, Pain level and pain-related behaviour classification using gru-based sparsely-connected rnns (2022).

12. A. Rényi, *Proceedings of the Fourth Berkeley Symposium on Mathematical Statistics and Probability, Volume 1: Contributions to the Theory of Statistics* (University of California Press, 1961), pp. 547–561.

13. C. Tsallis, *Journal of statistical physics* **52**, 479 (1988).

14. F. Kaspar, H. Schuster, *Physical Review A* **36**, 842 (1987).

15. T. Higuchi, *Physica D: Nonlinear Phenomena* **31**, 277 (1988).

16. H. Umezawa, H. Matsumoto, M. Tachiki, *Thermo Field Dynamics and condensed states* (North-Holland, Amsterdam,The Netherlands, 1982).

17. H. Umezawa, *Advanced field theory: Micro, macro, and thermal physics* (AIP, New York, U.S.A., 1993).

18. M. Blasone, P. Jizba, G. Vitiello, *Quantum Field Theory and its macroscopic manifestations* (Imperial College Press, London, U.K., 2011).

19. E. Celeghini, M. Rasetti, G. Vitiello, *Annals of Phys.* **215**, 156 (1992).

20. A. Chiolerio, M. Dehshibi, G. Vitiello, A. Adamatzky, *Symmetry* **14**, 1792 (2022).

21. A. Chiolerio, G. Vitiello, M. Dehshibi, A. Adamatzky, *Biomimetics* **8(1)**, 122 (2023).

22. A. Perelomov, *Generalized Coherent States and Their Applications* (Springer, Berlin, Germany, 1986).

23. C. Gerry, P. Knight, *Introductory Quantum Optics* (Cambridge University Press, Cambridge, U.K., 2005).

29. R. Hilborn, *Chaos and Nonlinear Dynamics* (Oxford University Press, Oxford, UK, 1994).

30. D. Bernal-Casas, G. Vitiello, *Barcelona-Salerno Preprint 2023* .
